# The Mp*ANT*-auxin loop modulates *Marchantia polymorpha* development

**DOI:** 10.1101/2024.12.03.626703

**Authors:** Melissa Dipp-Alvarez, J. Luis Lorenzo-Manzanarez, Eduardo Flores-Sandoval, Domingo Méndez-Álvarez, Annie Espinal-Centeno, Jesús León-Ruiz, Fernando Olvera-Martínez, John L. Bowman, Mario A. Arteaga-Vázquez, Alfredo Cruz-Ramírez

## Abstract

*AINTEGUMENTA-LIKE/PLETHORA* (*APB*) genes are considered part of the ancestral developmental toolkit in land plants. In *Arabidopsis thaliana*, these transcription factors are induced by auxin and are primarily expressed in tissues with actively dividing cells, where they play essential roles in organ development. *Marchantia polymorpha*, a liverwort that diverged from *A. thaliana* early in embryophyte evolution, possesses a single *APB* ortholog, Mp*AINTEGUMENTA* (Mp*ANT*), encoded in its genome. In this study, we aimed to characterize the function of Mp*ANT*. Analysis of a transcriptional fusion line indicates that Mp*ANT* is predominantly expressed in the meristematic region. We report that the Mp*ANT* promoter region contains several *cis*-acting Auxin Responsive Elements (AREs) and demonstrate that its expression, which occurs predominantly in meristematic regions, is significantly altered by addition of exogenous auxin and inhibition of auxin transport. These findings indicate that Mp*ANT* acts downstream of Auxin Response Factors (ARFs) and auxin signaling. Analyses of loss- and gain-of-function Mp*ANT* alleles highlight the importance of this transcription factor in meristem maintenance and cell proliferation. Additionally, we found that Mp*ANT* acts upstream of the auxin transporter Mp*PIN1* by influencing auxin distribution. Taken together, our findings reveal a feedforward regulatory loop involving auxin, Mp*ANT*, and Mp*PIN1* that is important for Marchantia development.

## 1 INTRODUCTION

The evolution of the land plant body, from the few cells of their zygnematal ancestors to the intricate three-dimensional structures of contemporary plants, was a key component in the radiation of land plants (Kenrick & Crane, 1997; Dolan, 2009; Donogue et al., 2020; Bowman, 2022). Recent studies have shown that members of early-diverging lineages of streptophytes share with derived angiosperms a set of conserved regulatory gene families and transcription factors (TFs). These TF families mainly evolved before land plant diversification, suggesting that a conserved core of regulatory genes was already present in the common ancestor of land plants (Catarino etal., 2016; Bowman et al., 2017; Lethi-Shiu et al., 2017; Wilhemsson et al., 2017; Hirakawa, 2022)

*AINTEGUMENTA-LIKE/PLETHORA/BABYBOOM* (APB) genes are considered one of these key developmental gene families in land plants (Floyd & Bowman, 2007). APB proteins contain two AP2 domains separated by a linker region of approximately 25 amino acids (Riechmann & Meyerowitz, 1998; Dipp-Alvarez & Cruz-Ramirez, 2019). In *Arabidopsis thaliana*, this family includes eight genes: *AINTEGUMENTA* (*ANT*), *AINTEGUMENTA-LIKE 1* (*AIL1*), *BABY BOOM* (*BBM*), and the *PLETHORAs* (*PLT1*, *PLT2*, *PLT3*, *PLT5*, *PLT7*), which are considered master regulators of diverse developmental processes (Reviewed in Horstman et al., 2014) and are mainly expressed in dividing tissues, where they regulate the maintenance of stem cell niches, the correct development of the embryo, and the formation of root and shoot organs (Aida et al., 2004; Galinha et al., 2007).

The liverwort *Marchantia polymorpha* L. ssp. *ruderalis* is a member of the bryophyte lineage and has benefited plant biology since the 18th century through its use as a model species for developmental, genetic, and physiological studies (Bowman et al., 2022; Kohchi et al., 2021). In addition to the molecular and genomic tools developed for this liverwort, *M. polymorpha* has low genetic redundancy due to the lack of whole-genome duplications (WGDs) (Bowman et al., 2017). These features make it an ideal model for studying the evolution of genetic systems that underlie major changes in land plant morphology and physiology (Delaux et al., 2019; Ishizaki, 2017; Bowman et al., 2022).

There is a single APB ortholog encoded in the genome of *M. polymorpha*, which we named Mp*ANT* (Mp8g11450, Dipp-Alvarez, M. & Cruz-Ramírez, A. 2019). Studies on *Arabidopsis* APB TFs have uncovered a relationship with the phytohormone auxin, which controls key aspects of plant development (Reviewed in Horstman et al., 2014). In *Marchantia*, Flores-Sandoval et al. (2018) explored TFs that co-express with hormonal signaling pathway genes at various developmental stages of the life cycle. In the 96-hour sporeling, a mild expression of Mp*ANT* was detected. Mp*ANT* belonged to a co-expression cluster with auxin signaling, biosynthesis and transport genes Mp*ARF1* (Flores-Sandoval et al 2016; Kato et al., 2017), Mp*ARF2* (Kato et al., 2020), Mp*IAA* (Kato et al., 2015; Flores-Sandoval et al 2015 ), Mp*YUC2* (Eklund et al., 2015l), Mp*PIN1* (Fisher et al., 2023), Mp*SHI*, and Mp*GRF*, indicating that it could be associated with in the auxin signaling pathway in *Marchantia*. Furthermore, Mp*ANT* expression is highly enriched in the apical cell tissue, similar to the expression observed in *A. thaliana* (Flores-Sandoval et al., 2018). This led us to investigate whether the single APB transcription factor Mp*ANT* could have a conserved function in meristem maintenance and plant development and, to evaluate the potential role of auxin signaling in its function. Auxin-mediated meristem establishment is partially controlled by long PIN intercellular auxin exporters in Marchantia, as Mp*pin*1^ge^ mutants are delayed in meristem formation relative to wild type in the sporeling to thallus developmental transition (Fisher et al., 2023). Two recently published findings parallel ours regarding the role of Mp*APB* in meristem maintenance during early gemma development (Fu et al., 2024; Liu, et al., 2024). However, here we provide novel evidence showing that Mp*ANT* forms a positive feedforward loop with auxin distribution and transport, defining not only its meristematic function but also its influence on the overall development of *Marchantia*.

## 2 METHODS

### 2.1 *Marchantia polymorpha* growth conditions and treatments

The Takaragaike-1 (Tak-1; Japanese male accession) was used as *M. polymorpha* wild-type in this study. The plants were cultured on half-strength Gamborg’s B5 (PhytoTechnology Laboratories) containing 0.5 g/L MES, 1% sucrose, and 1.3% phytoagar (PhytoTechnology Laboratories) (pH 5.7) and grown under continuous light (50∼60 μmol photons m⁻²s⁻1) at 22° C. For treatments, the gemmalings were grown on half-strength Gamborg’s B5 with 10 μM 2,4-D (Sigma-Aldrich), an auxin analogue (Flores-Sandoval et al., 2015) and 10 μM NPA (N-1-naphthylphthalamic acid), an inhibitor of directional (polar) transport of the hormone auxin (Abas et al., 2021).

### 2.2 Genetic constructs and *Agrobacterium*-mediated plant transformation

The MpANT coding sequence without stop codon was amplified from a pCRII plasmid generated by Eduardo Flores-Sandoval at John Bowman’s Laboratory (School of Biological Sciences, Monash University), with primers that contain attB1 and attB2 sequences for Gateway Cloning System. The MpANT (pMpANT) promoter fragment was cloned from *Marchantia polymorpha* Tak-1 DNA using primers designed with attB1 and attB2 tails for the Gateway Cloning System, the forward primer aligns with a sequence 2.5 kb upstream of MpANT start codon while the reverse primer hybridizes just before the start codon. Both sequences were amplified using polymerase chain reaction (PCR) with Accuprime High Fidelity polymerase. The *p*Mp*ANT:GFP* expression vector was constructed by LR-recombining the 2.5 kb *p*Mp*ANT* fragment in pKGWFS7 binary destination vector. The *pro35S:MpANT-CITRINE* expression vector for constitutive expression was constructed by recombining the CDS of MpANT without a stop codon in the pMpGWB206 binary destination vector. For gene editing via CRISPR-Cas9 of Mp*ANT*, a guide RNA (5’ GTTGGACGGGGCGCTACGAGG 3’) was designed to target a 21 bp sequence in the second intron - third exon boundary of the Mp*ANT* genomic region. The gRNA was cloned under the *MpU6* promoter into the GE010 plasmid (Sugano *et al.,* 2014), which also carries the Cas9 coding sequence. All plasmids were transformed into wild-type (WT) *M. polymorpha* plants of the Tak1 background through regenerating thalli transformation (Kubota et al., 2103). The regenerating plants weretransferred to half-strength Gamborg’s B5 (PhytoTechnology Laboratories) containing 0.5 g/L MES, 1% sucrose, 1.3% phytoagar (PhytoTechnology Laboratories) (pH 5.7), cefotaxime, and the appropriate selection antibiotics. Resistant transgenic plants from the CRISPR-Cas9 edition were genotyped and sequenced to confirm the nature of the edition. Two independent *p*Mp*ANT:GFP* transformant lines were obtained and analyzed for overall Mp*ANT* transcriptional pattern, while only *p*Mp*ANT:GFP-A* was used in auxin treatments. Two independent *pro35S:MpANT-CITRINE* lines with similar phenotype were obtained and further analyses were carried out with *pro35S:MpANT-CITRINE-A.* One CRISPR-Cas9 edited line showed phenotypic alterations and successful editions that resulted in a frameshift causing a premature stop codon which leads to the translation of a 415 AA protein instead of the 825 AA wild-type MpANT protein (Figure S1).

### 2.3 *In silico* genome-wide APB DNA-binding motif search

For the genome-wide APB DNA-binding motif search a Motif Discovery software for UNIX operating systems called HOMER (Heinz et al., 2010), was used. The tools seq2profile.pl and scanMotifGenomeWide.pl were used to make a motif file for the consensus APB DNA-binding sequence (CNTNGNNNNNNGTGC) reported by Santuari et al. (2016), and with this motif file, the whole *M. polymorpha* genome was scanned for motif occurrences. Once a motif coordinate output was generated, the motifs were annotated to a chromosome, the closest genome feature, and its gene ID, using UROPA (Kondili et al., 2017). The config file to run UROPA included instructions to only annotate motifs located up to 2500 bp upstream of a genome feature identified as exon or CDS. The output table contains the ID of the closest gene and was explored, filtered, and complemented with the *M. polymorpha* genome V6.1 (Iwasaki et al., 2021) information using the dplyr package in RStudio.

A Gene Ontology (GO) term analysis was conducted to find which biological processes are associated with the putative transcriptional targets of MpANT. For this, the list of genes with 1 or more MpANT binding sites was submitted to the GO Term Enrichment tool in PlantRegMap (http://plantregmap.gao-lab.org/go.php), selecting *M. polymorpha* as the input species. The p-value threshold was set to ≦0.05. The significantly enriched GO categories were visualized with the REVIGO web server (Supek et al., 2011).

A hierarchical clustering heatmap was generated using a list of putative MpANT targets related to auxin and development to see the TPM value of such genes in different *M. polymorpha* tissues. This analysis and heatmap were generated using the MBEX platform (<u>mbex.marchantia.info/clustergram</u>; Kawamura S., *et al*. 2022).

### 2.4 RT-qPCR methodology

Total RNA was extracted from wild type (Tak-1) and Mp*ant* thalli at 7 and 14 days post-sowing (dps) using the TRIzol method (Life Technologies). cDNA was synthesized in optimized conditions from the RNA samples with the SuperScript II (Thermo Fisher Scientific) using 2.5 μg of RNA in a 20 μL reaction volume utilizing Oligo dT (Sigma). Specific mRNA expression levels were determined by RT-qPCR utilizing the PCR SYBR Green master mix (Thermo Fisher Scientific) in a reaction volume of 20 μL. Three biological pools were used with three technical replicates per analyzed condition, the oligonucleotide efficiency was included in the calculations. Calculations were normalized with the expression level of the control gene Mp*ACT1*. The relative expression was calculated with the 2^(-ΔCт) method.

### 2.5 *In silico* analysis of MpPIN1 for the use of the antibody raised against AtPIN1

The antibody against AtPIN1 (aP-20, Santa Cruz Biotechnology, sc-27163) was designed to bind to the internal region of AtPIN1. To assess the potential of this antibody to bind the Marchantia ortholog, MpPIN1, the conservation of this region in MpPIN1 was evaluated. The sequence alignment and identity were determined using MUSCLE (https://www.ebi.ac.uk/jdispatcher/msa/muscle?stype=protein) with default settings. The hydrophilic loop region was defined using data from PDB 7Y9T (Yang *et al*., 2022), and the 2D structure of AtPIN1 was generated using Biotite (Kunzmann *et al*., 2023).

### 2.6 Immunohistofluorescence for IAA and MpPIN1 detection

The gemmae were fixed with 2.5% PFM in 1X PBS and Tween 20 at 4°C overnight, then washed three times with 1X PBS 0.2% Tween 20 at 4°C.Subsequently, the cell wall was degraded with 1% driselase (Sigma, D8037), 1% BSA (Sigma, A9647) at 37°C for 45 min and washed with 1X PBS 0.2% Tween 20 for 15 min. Posteriorly, the samples were permeated in 1X PBS 2% Tween 20 for 2 hours, on ice. For labeling, samples were incubated with primary polyclonal anti-IAA antibody (Agrisera, AS09 445) and polyclonal anti-PIN1 (Santa Cruz Biotechnology, aP-20: sc-27163) at a 1:1000 dilution, and incubated overnight at 4°C. Then washed three times with 1X PBS 0.2% Tween 20 for 6 hours at 4°C. For secondary labeling, the samples were incubated with anti-Rabbit IgG Alexa Fluor® 488 conjugate (Thermo Fisher Scientific, A11008) and anti-Goat IgG Alexa Fluor® 594 conjugate (Jackson, 305-586-045) at a 1:1000 dilution, and incubated overnight at 4°C. Then washed three times with 1X PBS 0.2% Tween 20, on ice. Afterwards, the samples were treated with DAPI (Sigma, D9542) to label nuclei. Finally, samples were washed with 1X PBS 0.2% Tween 20 for 1 hours and mounted in a coverslip with Prolong-Gold antifade mountant reagent. Finally, samples were washed with 1X PBS 0.2% Tween 20 for 1 hours and mounted in a coverslip with Prolong-Gold antifade mountant reagent. Conditions for IAA immunolabeling were adjusted based on Aguilar-Cruz et al., (2023).

### 2.7 Tissue staining and Imaging

The gemmae and thalli were stained with Propidium Iodide (PI) at a concentration of 10 mg/L in ddH2O for 10 minutes in an Eppendorf tube. After incubation, the samples were rinsed twice with ddH2O. EdU staining was carried out similarly to Fu et. al., 2024, thalli of 4 days-post-sowing were immersed in 1/2 strength Gamborg’s B5 liquid medium with 10 μM EdU at 22°C for 2 hours under white light. Then they were fixed in FAA (50% Ethanol, 2.5% glacial acetic acid, 2.5% formaldehyde) for 2 hours at room temperature. Then samples were washed with PBST buffer (0.5% Triton X-100, pH 7.4) for 5 minutes and PBS buffer (pH 7.4) for 5 minutes. Then the samples were labeled with Yefluor 488 Azide (40287ES60, YEASEN) for 30 minutes in dark conditions. After the samples were washed with PBS buffer (pH 7.4) for 30 min.

### 2.8 Keyence microscopy

Gemmae and thalli of 0, 4, 14 and 30 days-post-sowing were observed using a Keyence VHX-5000 Digital Microscope (Keyence Corp.) with an LED light source and a 20-200X magnification range lens (VH-Z200).

### 2.9 Confocal microscopy

Confocal laser scanning microscopy imaging of gemmae, apical notches, rhizoid initial cells and thalli was performed on an inverted LSM-800 Zeiss microscope (Zeiss, Oberkochem, Germany). Propidium iodide was excited at 561 nm with a laser intensity of 0.2%, emitted light was collected between 576 nm and 700 nm. eGFP was excited at 488 nm with a laser intensity of 20%, emitted light was collected between 410 nm and 546 nm. For immunolocalization, the IAA signal was excited at 488 nm with an emission wavelength of 520 nm. The PIN1 signal was excited at 590 nm, and the emission wavelength was 617 nm. DAPI staining signal was detected with an excitation wavelength of 405 nm and emission wavelength of 461 nm.

### 2.10 Image processing

The area (mm^2^) of gemmae and thalli were measured using the Measure tool of Fiji (https://imagej.net/software/fiji/). To measure the number of rhizoid initial cells, images of gemmae stained with PI were analyzed using Fiji. The Z-stack immunolocalization images were processed in Fiji using the Z-project function with maximum intensity projection. For nuclei analysis, the Mean-Shifh Super Resolution (MSSR) method (Torres-García et at., 2022) was applied with the following parameters: AMP= 5, FWHM of PSF= 4, and Order=1. To analyze the results, ANOVA was performed followed by a Post-Hoc Tukey HSD test.

## 3 RESULTS

### 3.1 Auxin positively modulates Mp*ANT* expression in the meristematic region of *Marchantia polymorpha*

The protein structure of MpANT is similar to that of *A. thaliana* APB proteins, and the sequence and length of the DNA-binding AP2-R1 – linker – AP2-R2 region display a high degree of conservation (FIGURE 1A-B). The spatiotemporal transcriptional pattern of Mp*ANT* was characterized using a construct containing a 2.5 kb sequence upstream of the Mp*ANT* transcription start site (TSS) fused to the GFP gene (*_pro_*Mp*ANT*:*GFP*). GFP signal was observed in the apical notch regions (Figure 1C, mock) of young gemmae. Previous studies have shown that Mp*ANT* is co-expressed with Mp*ARF1* and Mp*ARF2* in the apical meristem region of *M. polymorpha* (Flores-Sandoval et al., 2018). Consistent with a putative link between MpARFs and Mp*ANT*, we identified 6 putative auxin response elements (AREs) in the Mp*ANT* -*cis* regulatory region (Figure 1B; Dipp-Álvarez & Cruz-Ramirez, 2019).

To investigate the role of auxin in regulating Mp*ANT* transcription, we plated gemma in mock, the auxin analogue 2,4-D (10μM), the auxin transport inhibitor NPA (10μM), or a combination of NPA+2,4-D and observed the expression driven by *_pro_*Mp*ANT*:*GFP* 4 days post plating (dpp). The application of 2,4-D expanded the GFP expression domain, with signal detected not only in the apical notch but also in the transition zone (Figure 1C). Inhibition of auxin transport with NPA resulted in increased signal at the apical notch, suggesting that Mp*ANT* expression is influenced by auxin accumulation. Furthermore, the combination of exogenous auxin and transport inhibition expanded the expression pattern and created additional auxin maxima, as observed in the NPA+2,4-D-treated plants (Figure 1C). Together, these results indicate that auxin signaling, perhaps through the MpARF1 activator, modulates Mp*ANT* transcription in response to auxin levels and distribution.

**FIGURE 1.**
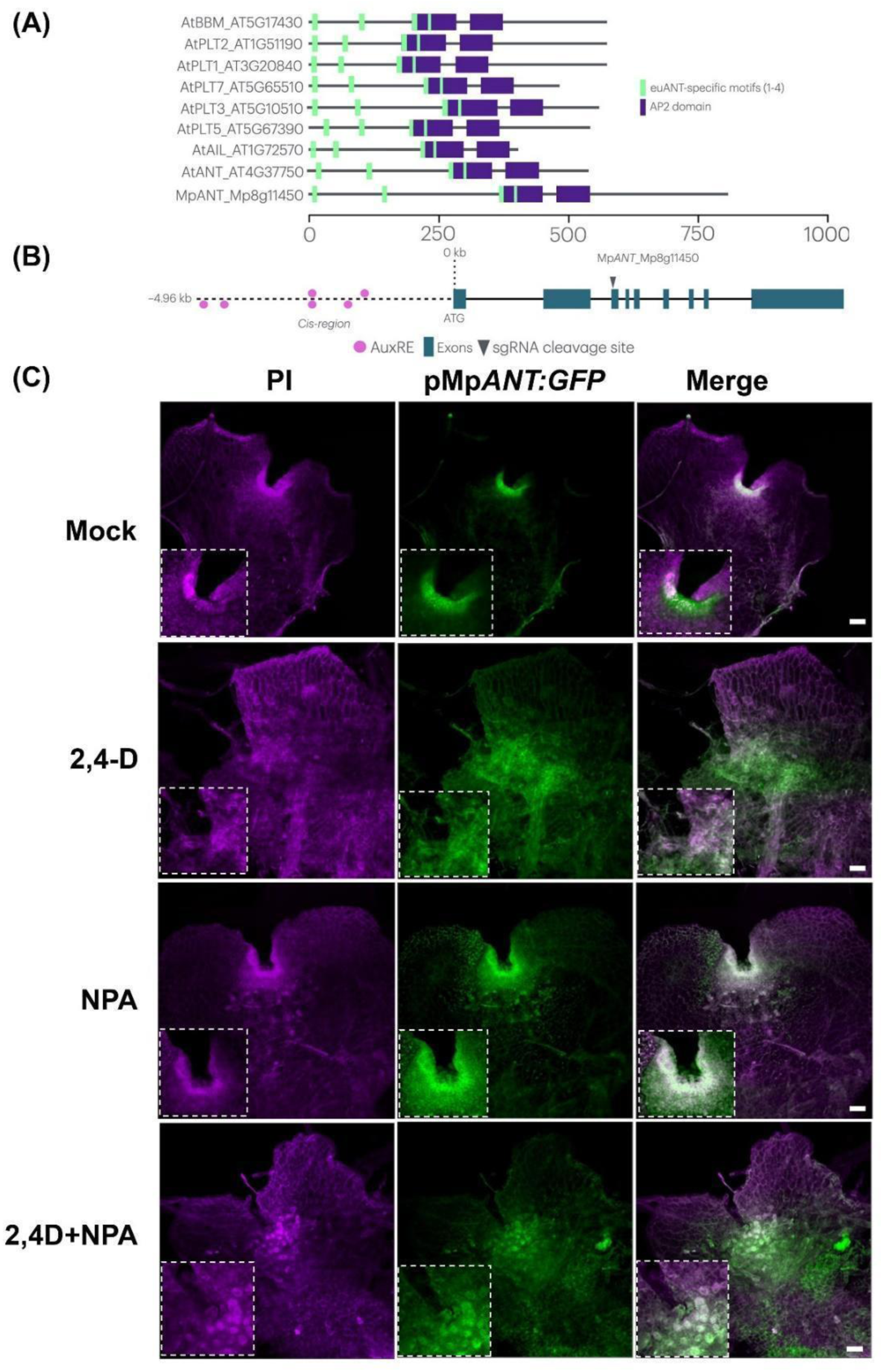
Mp*ANT* transcription is responsive to fluctuating auxin. (A) Conserved domain organization and euANT-specific motif of *A. thaliana* and *M. polymorpha* euANT clade members. (B) Gene model for Mp*ANT* with AREs elements demarcated in its promoter region (pink dots) and the target region for editing with CRISPR-Cas9 (arrow head). (C) Expression analysis of *_pro_*Mp*ANT:GFP M. polymorpha* line. Confocal images show the expression patterns of 4-day-old wild-type gemmae grown in Gamborg’s B5 media (mock), or gamborg media supplemented with either 2,4-D (10μM), NPA (10μM) and NPA+2,4-D. Green fluorescence corresponds to GFP signal, purple fluorescence corresponds to Propidium Iodine signal. Insets in all panels are magnifications of the apical notch. Scale bars=100 μm.

### 3.2 Gain- and loss-of-function alleles indicate Mp*ANT* plays key roles in *M. polymorpha* early development

To investigate the biological roles of Mp*ANT* in *Marchantia polymorpha* early gemma development, both gain- and loss-of-function lines were generated. For the gain-of-function analysis, we generated an Mp*ANT* ectopic overexpression line with a translationally fused fluorescent reporter (*_pro_35S:*Mp*ANT-CITRINE*; hereafter referred to as Mp*ANT^OE^*). To generate Mp*ant* loss-of-function mutant alleles (hereafter referred to as Mp*ant*) were generated as described in the Methods section and Figure S1.

We observed that Mp*ant* gemmae were smaller than WT, with area quantification supporting a statistically significant reduction in size (Figure 2A and D). In contrast, Mp*ANT^OE^* gemmae were significantly larger than WT and Mp*ant* plants (Figure 2A and E). These size differences were also evident in Mp*ant*, Mp*ANT^OE^*, and WT young thalli, not only in the overall area but also in the putative meristematic region (Figure 2B and E).

To correctly assess the impact of Mp*ANT* on cell proliferation in the meristematic regions, we performed EdU staining experiments on Mp*ant*, Mp*ANT^OE^*, and WT thalli. We observed a reduction in the proliferative cell area in Mp*ant* thalli compared to WT controls (FIGURE 2C and F). This finding aligns with recent studies characterizing loss-of-function mutants of Mp*ANT*, although in different *Marchantia* strains (Fu et al., 2024; Liu et al., 2024). In all three independent loss-of-function mutants, downregulation or absence of Mp*ANT* was found to compromise meristem function. In contrast, our results with Mp*ANT^OE^* plants revealed that Mp*ANT* ectopic overexpression expands the area of proliferative cells in young thalli, compared to WT plants, and even more so than in the loss-of-function mutants (Figure 2C and F).

We also found that Mp*ANT* function is important for rhizoid cell development. PI staining and confocal microscopy imaging of 2 days post-plating (dpp) gemmae from Mp*ant*, Mp*ANT^OE^*, and WT plants revealed that Mp*ant* gemmae developed fewer rhizoid initial cells compared to WT plants, while Mp*ANT^OE^* gemmae exhibited a greater number of rhizoid initials (Figure S2 A-D).

**FIGURE 2.**
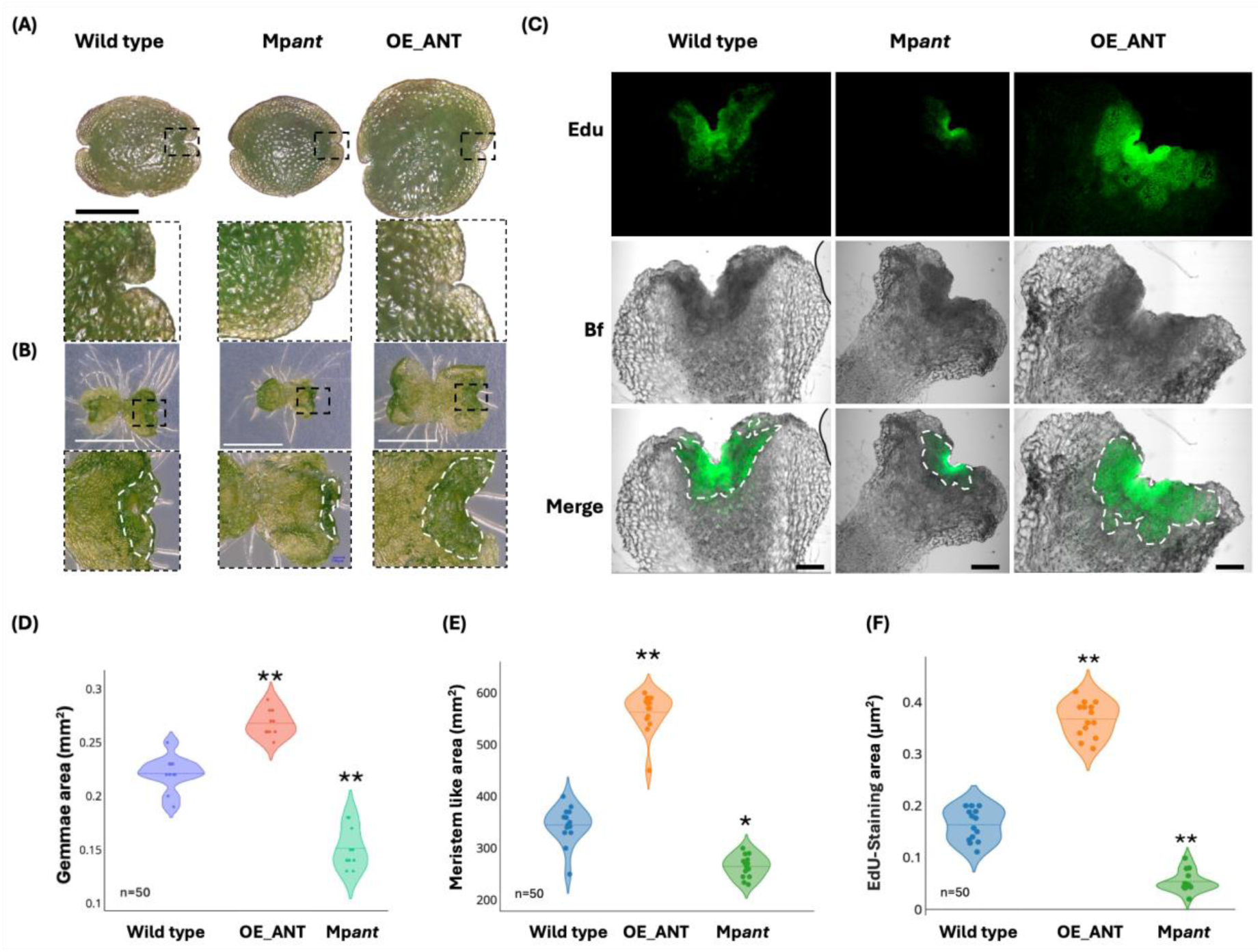
Mp*ANT* is important for meristem function in *Marchantia* thalli. Overall morphology of WT, Mp*ant* and Mp*ANT^OE^* thalli: (A) Gemmae morphology, the black dotted lines show a magnification of the apical notch. Black scale bar=0.2mm. (B) Thalli of 4 days-post-plated, the black dotted lines show the apical notches and white dotted lines show a magnification of the transition zones. White scale bar=2mm. (C) EdU incorporation assay to visualize the proliferative cells in the apical notch and transition zone of 4-day-old thalli. White dotted lines show Edu-stained cells. BF: Bright field. Edu scale bar=200 μm. (D) Gemmae area (mm2) of WT, Mp*ANT^OE^* and Mp*ant* (E) Meristem-like areas (mm2) of WT, Mp*ANT^OE^* and Mp*ant* young thalli. (F) Area of Edu-positive cells (μm2) in WT, Mp*ANT^OE^* and Mp*ant* young thalli. One-way ANOVA followed by Tukey’s post hoc test compared to WT (*P < 0.05 and **P < 0.01).

### 3.3 Mp*ANT* regulates thallus growth, branching architecture, and antagonizes gemma cup production in *M. polymorpha*

In *M. polymorpha*, thallus growth and organization are determined by the highly regulated division of apical and sub-apical cells in the apical notch (Shimamura, 2016). Our data show that Mp*ANT* plays a key role in meristem maintenance, influencing not only thalli growth and branching architecture which is also known to be auxin-dependent (Streubel et al., 2023). To further investigate the role of Mp*ANT*, we analyzed the branching phenotypes and gemma cup production in Mp*ant* and Mp*ANT^OE^* lines and compared them to WT plants.

At 14 dpp, Mp*ant* and Mp*ANT^OE^* lines exhibited distinct differences in branching patterns compared to WT plants (Figure 3A). The area of M*pant* lines was smaller than that of WT controls, while Mp*ANT^OE^* lines showed a statistically significant increase in size (Figure 3C). In the meristematic zone, WT and Mp*ANT^OE^* lines exhibited the typical convex shape at the apical notch and formed dichotomous branching points (Solly et al., 2017). In contrast, Mp*an*t lines showed delayed development, characterized by the formation of central lobes (Zoom-in, Figure 3A). All lines contained approximately twelve gemma cups at 14 days, consistent with previous studies (Susuki et al., 2020). At one month, significant differences in development area were observed among the lines (Figure 3B), though the number of plastochrons remained similar (Figure 3E). Notably, Mp*ant* lines exhibited a dramatic overproduction of gemma cups, approximately 70 per plant, compared to around 40 and 30 gemma cups in Mp*ANT^OE^* and WT lines, respectively (Figure 3F).

Since auxin influences Mp*ANT* expression (Figure 1), we examined whether the gemmae cup phenotype observed in Mp*ANT* loss- and gain-of-function mutants is related to auxin. We treated Mp*ant* and Mp*ANT^OE^*plants with NPA and exogenous auxin (2,4-D) and noted gemmae cup overproduction in 21 old Mp*ant* plants was drastically reduced from more than 30 gemmae cups in the mock medium to half of that in 2,4-D treated plants and to an average of 3 in plants grown in NPA-supplemented medium (Figure S3A and B). The effect of exogenous auxin and NPA on gemmae cup formation was even more pronounced in WT and Mp*ANT^OE^* plants which exhibited a complete inhibition in both treatments compared to its respective control (Figure S3 and B). These results together suggest that Mp*ANT* plays a negative role on gemmae cup formation. Although the negative role of auxin on gemmae cup formation has been previously described (Flores-Sandoval et al., 2015; Komatsu et al., 2023), our findings implicate Mp*ANT* in this process.

**FIGURE 3.**
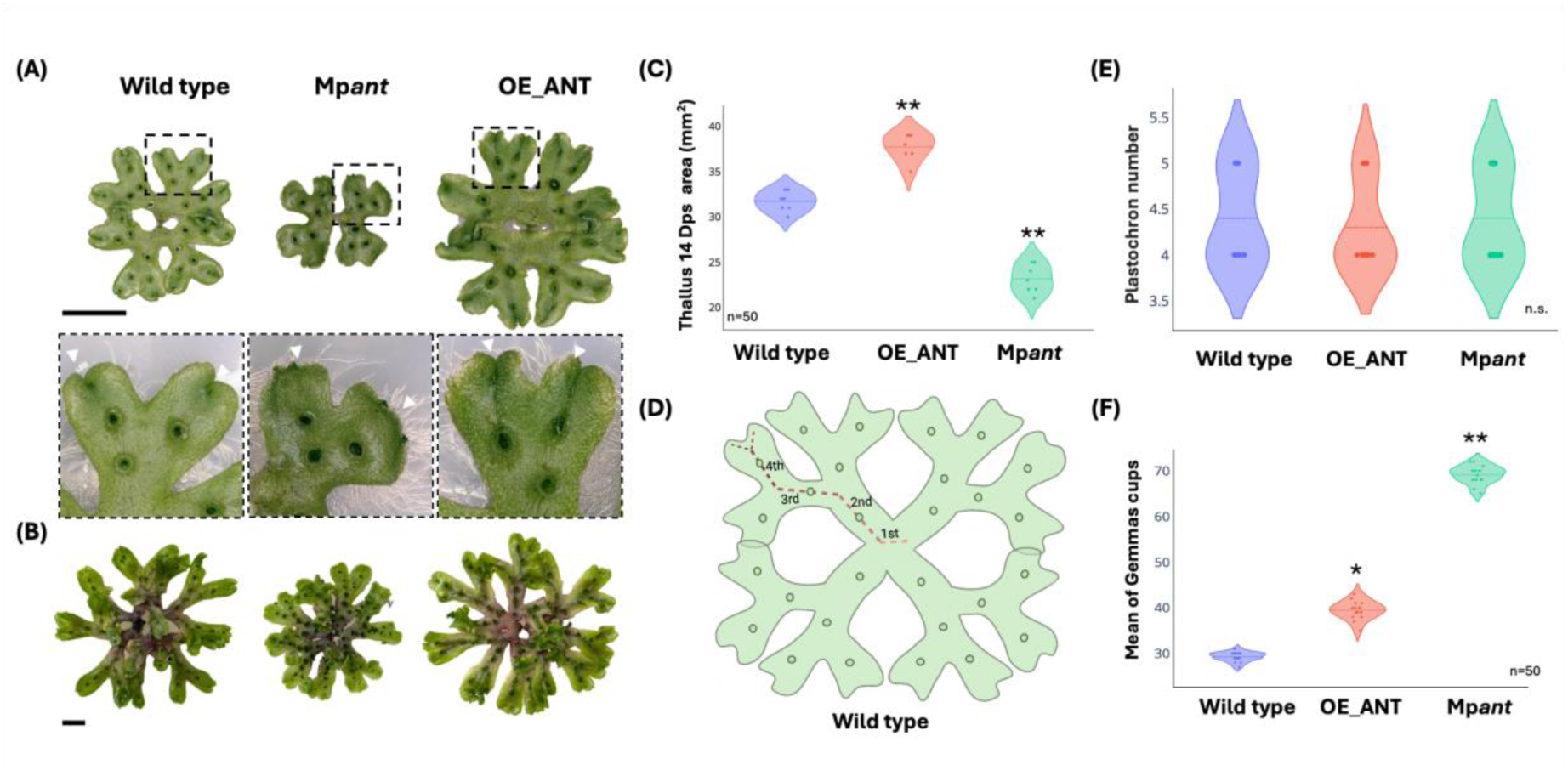
Plastochron and branching analyses of WT, Mp*ant* and Mp*ANT^OE^* lines. (A)Thalli of 14 dpp. The black dotted lines show zoom in meristem zone and the white arrowheads show differences in the notch divergence. (B) Phenotype of a 1-month-old WT, Mp*ant* and Mp*ANT^OE^* plants. (C) Area (mm2) of thallus 14 dpp (D) Normal plastochron and gemmae cup development in 1-month-old plants. (E) Plastochron numbers analysis in WT, M*pant* and Mp*ANT^OE^*plants at 30 days. (F) Mean of gemmae cup number at 30 days. Scale bars = 1 cm. One-way ANOVA followed by Tukey’s post hoc test compared to Tak-1(*P < 0.05 and **P < 0.01), n.s.=non-significant.

### 3.4 Genome-Wide identification of MpANT binding sites reveals its role in development and auxin signaling

To identify potential targets of MpANT, we conducted a genome-wide search in the *M. polymorpha* genome using the consensus APB DNA-binding motif (CNTNGNNNNNNGTGC), as reported by Santuari et al. (2016). Our results show that approximately 13% (2515 genes) of the protein-coding genes in the *M. polymorpha* genome could be regulated by MpANT (Figure S4A). Most of these predicted target genes harbor a single APB binding site, and the majority are located within 500 bp upstream of their transcription start sites (Figure S4B). Gene ontology analysis of the predicted target genes revealed that many are involved in critical biological processes such as development, morphogenesis, hormone signaling, and triterpene biosynthesis. Notably, a recent study by Wu et al. (2024), which performed ChIP-Seq to identify Mp*ANT* targets, found overlap with our predictions, as 387 genes were shared between both datasets (Table S1).

Given that the Mp*ANT* promoter is responsive to different auxin levels, we further examined whether auxin-related genes are regulated by MpANT. Our analysis revealed that the APB binding site is present in the promoter regions of Mp*PIN1* and Mp*TAA*, two genes involved in auxin transport and synthesis, respectively. These genes are co-expressed in the apical cell region, where Mp*ANT* expression is also prominent (Figure S5). We also performed a co-expression analysis to identify genes whose transcript levels are highly correlated with Mp*ANT* expression across various tissues and developmental stages. Among the putative MpANT targets, we identified genes whose orthologs in *A. thaliana* are critical for development. Notably, Mp*MADS2*, a MIKCc-type MADS-box gene, was found to be expressed at higher levels in gametangia (Figure S5) and is a predicted target of MpANT. In *A. thaliana*, there is evidence for interaction between the APB family and MIKCc-type genes such as *AGAMOUS* (Krizek et al., 2020). Another putative Mp*ANT* target, Mp*SPL2* (Mp1g10030), is homologous to the SQUAMOSA PROMOTER-BINDING LIKE (SPL) TFs in *A. thaliana*, which regulate the transition to the reproductive phase and the development of floral organs.

These findings indicate that MpANT regulates diverse processes in *M. polymorpha* by modulating the expression of genes involved in development, auxin signaling, and stress responses. To explore the role of MpANT in auxin-related pathways, we analyzed the transcript levels of several auxin-related genes in Mp*ant* lines at 7 and 14 days post-germination (dpg) by qRT-PCR. At 7 dpg, genes such as Mp*YUC1*, Mp*YUC2*, Mp*ARF1*, Mp*ARF3*, Mp*TAA*, and multiple Mp*PIN* genes were moderately downregulated in Mp*ant* plants compared to WT controls (Figure 4A). At 14 dpp a stronger downregulation of Mp*ANT*, Mp*ARF1*, Mp*ARF2*, Mp*ARF3*, Mp*TAA*, Mp*PIN1* and Mp*PIN2* was observed (Figure 4B).

Additionally, we examined the expression of Mp*GRAS3* (Mp1g10440) and Mp*CYCD*, which are involved in cell proliferation and meristem maintenance in Arabidopsis root stem cell niche. Both genes showed slight downregulation in Mp*ant* plants compared to the control (Figure 4B).

**FIGURE 4.**
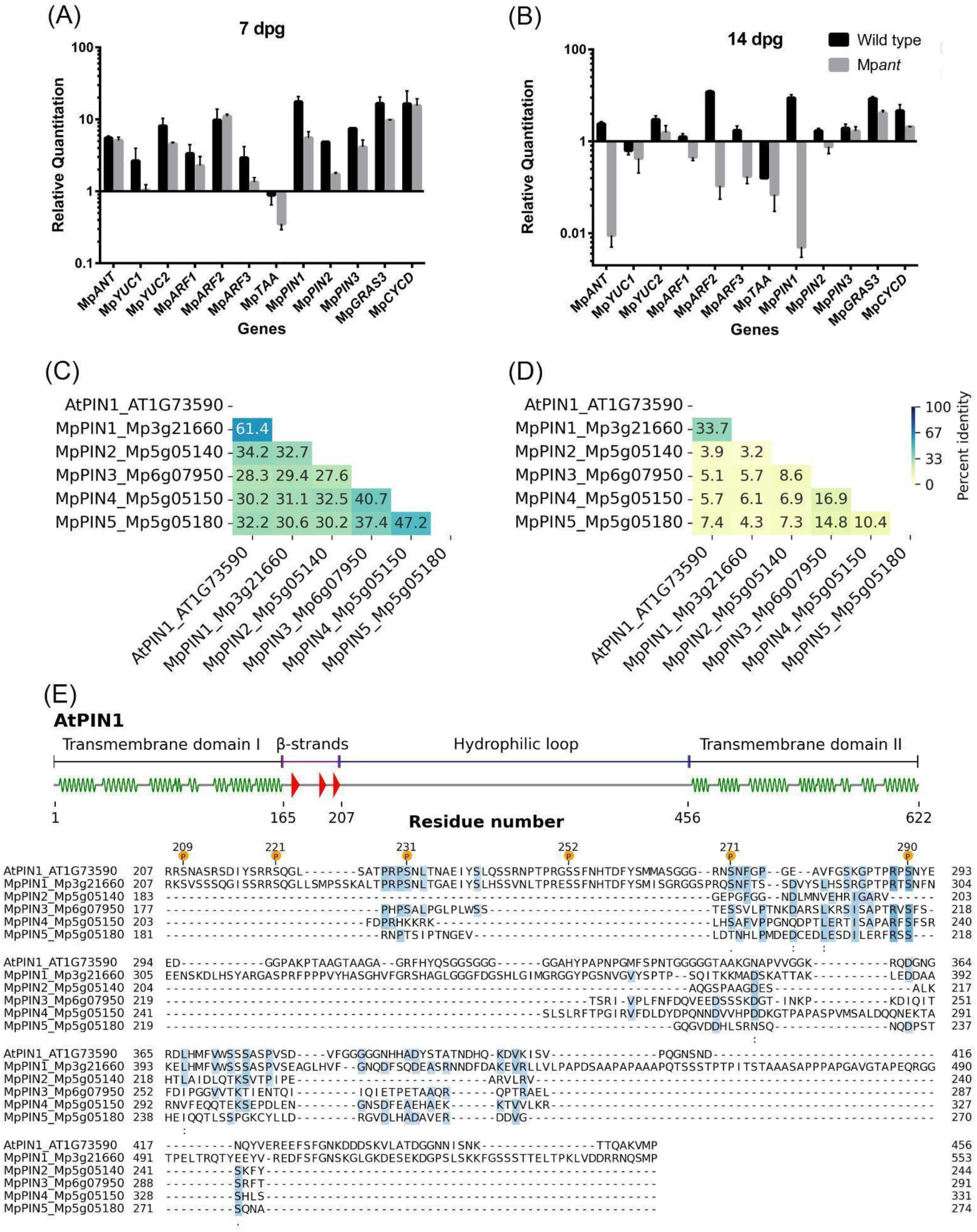
MpANT regulates the transcription of genes involved in auxin synthesis and auxin transport, including Mp*PIN1* which is the sole ortholog of long AtPINs. qRT-PCR analyses of auxin-related genes and putative MpANT targets at 7 (A) and 14 dpg (B) in WT and Mp*ant* plants. (C) Global percent identity of AtPIN1 against MpPINs. (D) Conservation of internal region in MpPIN1 antibody raised for AtPIN1. (E) Conservation of AtPINOID phosphorylation sites in the MpPIN1 hydrophilic loop (Fisher et al., 2023).

### 3.5 Mp*ANT* acts upstream of MpPIN1 localization and auxin distribution

We demonstrated that auxin concentration and distribution are crucial for proper MpANT expression in the apical notch. These findings, alongside the presence of several auxin response elements (AREs) in the Mp*ANT* promoter, suggest that MpARFs regulate Mp*ANT* expression in response to the auxin maxima formed at the apical notch. Previous studies (Flores-Sandoval et al., 2015) showed that disrupting auxin signaling affects gemmae cup formation and thallus branching, phenotypic alterations we observed in Mp*ant* loss- and gain-of-function lines. In Arabidopsis, *PLT* genes act downstream of auxin signaling but also regulate genes involved in auxin synthesis and transport in a feedback loop (Horstman et al., 2014).

Since Mp*ANT* acts upstream of Mp*PIN1* and Mp*PIN2*, we investigated the patterns of IAA distribution and MpPIN1 localization in WT, Mp*ant* and Mp*ANT^OE^* gemmaelings. For this we performed immunofluorescence assays with antibodies against IAA and MpPIN1. To assess the potential usefulness and specificity of a commercial antibody generated against AtPIN1 to recognize MpPIN1, we performed diverse *in silico* analyses (see methods) which indicated that AtPIN1 shares greater identity with MpPIN1 than with other *M. polymorpha* PIN proteins (Fisher et al 2023; Figure 4C-E and Figure S6), supporting the use of this antibody for specifically detecting MpPIN1. Furthermore, we corroborated that the phosphorylation sites critical for proper AtPIN protein localization are well conserved in MpPIN1 (Fisher et al 2023; Figure 4E).

Double immunofluorescence assays revealed significant differences in the localization of IAA and MpPIN1. In WT gemmae, auxin maxima were observed in the apical notch, rhizoid initials (green fluorescent signal, Figure 5A and B), and the gemmae stalk. MpPIN1 was also localized to the membranes of apical notch and rhizoid cells (red fluorescent signal, Figure 5A-B). However, in Mp*ant* gemmae, this pattern was drastically altered. MpPIN1 no longer localized to the cell membranes but instead re-localized to other organelle membranes, as observed in both the apical notch cells and rhizoid initials (Figure 5A-B). The mislocalization of MpPIN1 correlated with changes in IAA distribution, with the auxin signal shifting from the apical notch to the cytoplasm and membranes of rhizoid cells (Figure 5B).

**FIGURE 5.**
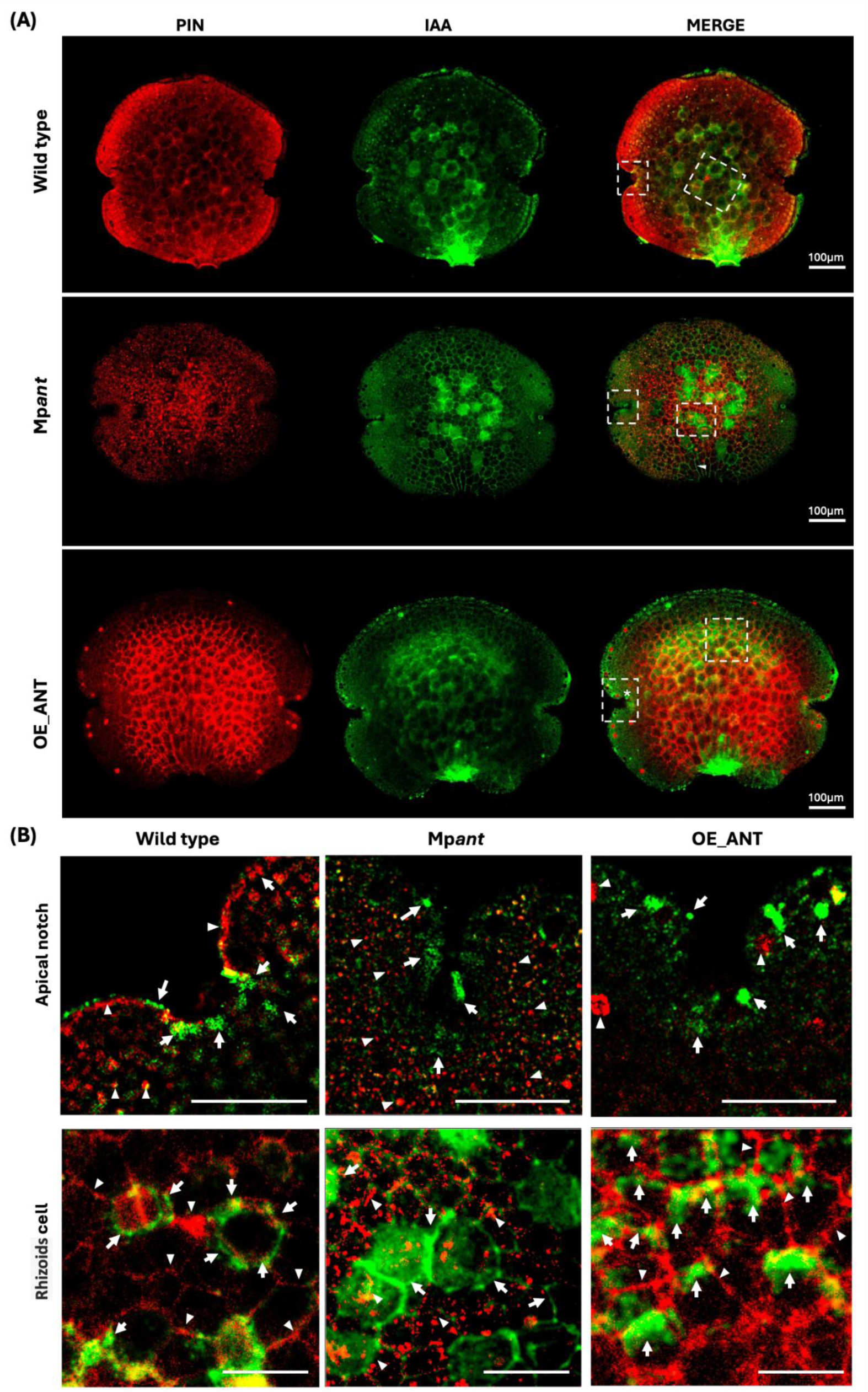
Mp*ANT* is important for polar localization of MpPIN1 and auxin distribution in *Marchantia* gemmae. Immunolocalization of MpPIN1 and IAA in Marchantia gemmae. (A) Confocal microscopy images of WT, Mp*ant* and Mp*ANT^OE^*. The red signal corresponds to PIN (Alexa Fluor 594 conjugated secondary antibody) and green signal to free IAA immunofluorescence (Alexa Fluor 488 conjugated secondary antibody). The white dotted lines show zoom in meristem zone and epidermal cells. Scale bars = 100 μm. (B) PIN and IAA distributions in apical notch and epidermal cells. In the notch zone, the white arrow shows IAA accumulation in the border cell, transitions zone, apical and sub apical cell, respectively. In epidermal cells, the white arrows show the IAA accumulation in the endomembrane and inside epidermal cells. In the notch area, arrowheads show PIN localization, accumulation in the border cell, peripheral zone, transition zone and oil body cells. In epidermal cells of WT and Mp*ANT^OE^* the arrowheads show the PIN accumulation in the endomembrane, while in Mp*ant* the PIN is localized also in the cytosol . Scale bars = 50 μm arrowhead.

Intrigued by the mislocalization of MpPIN1 in the Mp*ant* gemmae, we conducted further double immunofluorescence assays for MpPIN1 and auxin detection, staining with DAPI to denote the nuclei. In some cases, we observed co-localization of MpPIN1 with the DAPI signal, suggesting potential nuclear localization, as suggested by Tang et al. (2024). A more detailed analysis using Mean Shift Super-Resolution (MSSR) on WT, Mp*ant*, and Mp*ANT^OE^* gemmae revealed that MpPIN1 was localized outside the nucleus (Figure S7).

Immunolocalization assays showed differences in the distribution and accumulation of MpPIN1 and IAA signal in Wt, Mp*ant*, and Mp*ANT^OE^* gemmae (Figure 5). In WT, IAA and DAPI signals overlapped, while MpPIN1 was localized around the nucleus (Figure 6A and Figure S7). In Mp*ant* gemmae, IAA and MpPIN1 signals were also localized to the cytoplasm, far from the nucleus (Figure 6B). In contrast, Mp*ANT^OE^*lines exhibited highly polarized distribution of MpPIN1 at the plasma membrane, while IAA signal accumulated in the cytoplasm, and around the nuclei, with both signals co-localizing near the nuclei (Figure 6C). This is consistent with recent findings showing that overexpressed MpPIN1-GFP is polarized to the tips of young rhizoids, whereas overexpressed AtPIN1-GFP shows weak membrane localization and lacks polarity (Tang et al., 2024).

Our *in silico* comparisons among AtPINs and MpPIN1 proteins show that the phosphosites in AtPIN1, which are targets of AtPINOID, for the localization of AtPIN1 to the plasma membrane are highly conserved in MpPIN1, suggesting the existence of a AtPINOID (AtPID), and AGC kinase, homologs in *Marchantia*., and three homologs were identified in *Marchantia* (Bowman et al 2017; Figure S8A).

Moreover, since in the Mp*ant* gemmae the localization of MpPIN1 is drastically altered, we speculated whether MpANT could be influencing, indirectly, MpPIN1 localization by regulating the transcription of the putative MpAGCs*. In silico* analyses of the promoters of the MpAGCs show that in a 2 kb region upstream the ATG, all three MpAGC genes have several MpANT response elements (MpANT-REs) as well as several ARF AREs (Figure S8B).

**FIGURE 6.**
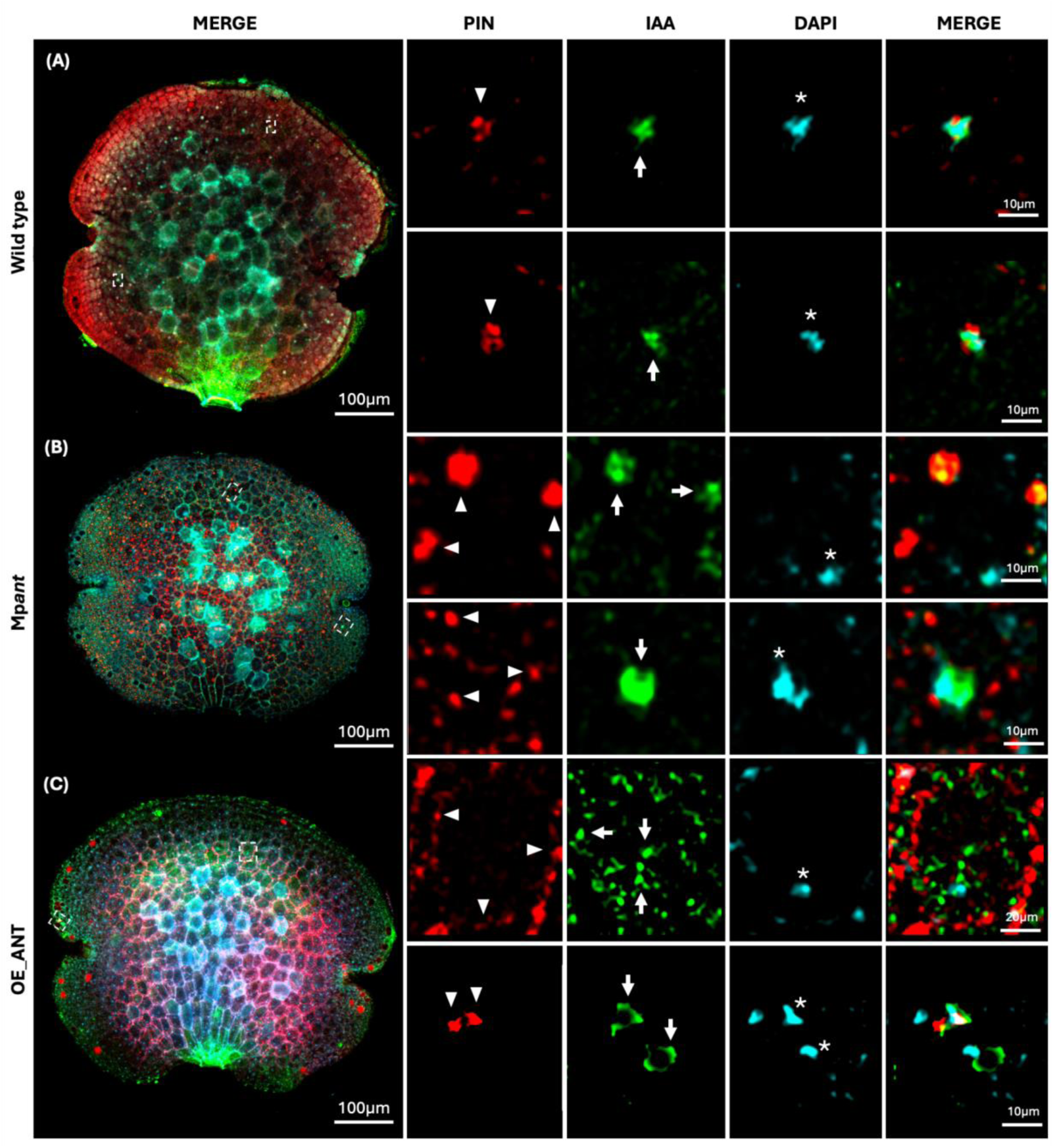
MpPIN1 is localized around the nucleus in Mp*ant* cells. Nuclear localization analyses of MpPIN1 and IAA in WT, Mp*ant* and Mp*ANT^OE^* gemma. The red signal corresponds to MpPIN1 (Alexa Fluor 594 conjugated secondary antibody), the green signal to free IAA (Alexa Fluor 488 conjugated secondary antibody), and cyan signal to DAPI (4’,6-diamino-2-phenylindole) marking nuclei. Arrowheads show the MpPIN1 signal, white arrows show the IAA accumulation and the asterisks depict nuclear localization. (A) In Wt, the two replicates show that the MpPIN1 signal surrounds IAA and nuclei signals. (B) The MpPIN1 signal is localized inside the cell, while IAA accumulation is present in the nuclei and together with MpPIN1 signals. c) The PIN1 signal was detected in the cell membrane and around nuclei signals, while the IAA signal was localized inside the cell and around nuclei.

**FIGURE 7.**
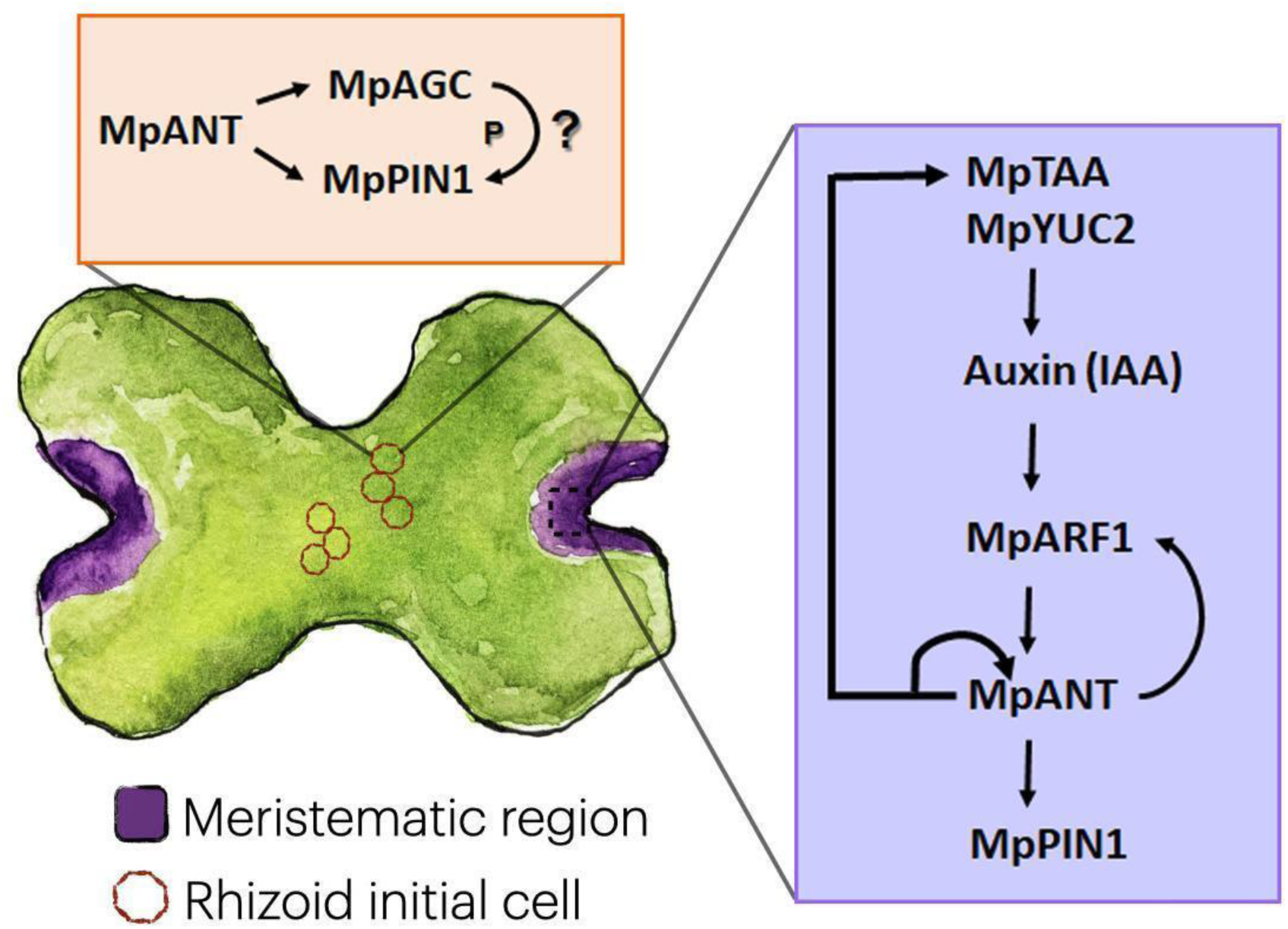
The roles of Mp*ANT* roles in regulatory networks in the meristem (purple box) and in epidermal cells (pink box) of Marchantia. Diverse studies and ours show the expression of Mp*ANT* in the apical notch, coincident with an auxin maximum, here we show that increasing auxin concentrations induces Mp*ANT* transcription, most probably via ARFs action. Mp*ANT* acts upstream of auxin synthesis genes Mp*TAA* and Mp*YUC2* as well as on Mp*PIN1* involved in auxin transport. This loop is essential for the proper development of the *Marchantia* meristem. Additionally, in the epidermal cells the localization of MpPIN1 to the plasma membrane depends on Mp*ANT*, probably by regulating the expression of putative AGC Kinases encoded in the *Marchantia* genome. The described functions for Mp*ANT* in the meristem and epidermal cells are not mutually excluded.

## 4 DISCUSION

### 4.1 A genetic program driven by Mp*ANT* maintains a stem cell niche in coordination with the auxin pathway

The PLETHORA TFs are essential for the maintenance of the stem cell niche in the root apical meristem of *A. thaliana.* PLTs act in gradients of expression and are key players of a network that establishes a feed forward positive loop for auxin maxima formation. PLT TFs directly regulate genes involved in auxin synthesis and transport, and conversely PLTs expression in the RAM depends on auxin maxima formation. Our study and two other recent ones (Fu et al., 2024; Liu et al., 2024) show that Mp*ANT* transcription is active in the meristem of *Marchantia* thalli and concur that loss-of-function Mp*ANT* mutants result in reduced meristematic activity leading to smaller plants (Figure 2). Therefore, Mp*ANT* is essential for the proper size and function of the stem cell niche in *Marchantia*. On the other hand, Mp*ANT* overexpression expands the region of proliferative cells in a pattern which suggests a correlation between auxin gradient and zonation of stem cells, its transit amplifying daughters and the final differentiation towards the midrib. Liu et al., (2024) proposed that Mp*ANT* restricts cell proliferation and meristem size through its transcriptional target Mp*CLE1*, however our EdU staining results show that overexpression of Mp*ANT* expands the meristem and the stem cell pool. We should explore whether any aspects of the Mp*ant* phenotype are Mp*CLE1*-dependent in future studies. Fu et al 2024 also propose that MpANT promotes meristem maintenance via MpWOX but Hirakawa et al., (2020) showed that Mp*wox* mutants are not meristem deficient. We should also consider that while we generated all our research in the widely distributed strain Tak-1, Liu et al, (2024) used the Upp strain.

### 4.2 A feed forward loop involving Mp*ANT*, auxin transport and auxin distribution modulates Marchantia development

The Mp*ANT* transcription domain is defined by auxin transport, distribution and concentration, most probably by the action of MpARFs on Mp*ANT* (Figure 1, Figure7), in turn MpANT induces the expression of its direct, and indirect, transcriptional targets, some of them related to auxin synthesis and auxin transport (Figure 4, Figure7). This network creates a positive loop for the meristem function in which MpANT modulates Mp*ANT* transcription (Figure7). Liu et al., (2024) position MpANT directly regulating Mp*GRAS9 ,* while Fu et al., (2024) proposed that MpANT regulates Mp*WOX.* Mp*GRAS3* and Mp*CYCD* are downstream MpANT (Figure 4). These observations indicate that the link between the two main pathways for stem cell niche maintenance in Arabidopsis, the Aux-AtPLT and the SHR-SCR-CYCD-RBR regulatory networks seem to be conserved in *Marchantia*. However, a clear definition of Marchantiás functional orthologs for several genes of these networks still needs to be defined.

Besides the auxin-related putative targets of MpANT, we found several genes highly co-expressed with Mp*ANT* such as Mp*PYL1*, the closest gene to Mp*ANT* in terms of co-expression, and a putative MpANT target. Mp*PYL1* encodes an ABA receptor protein, with orthologs in Arabidopsis (At*PYL4-6*) involved in the ABA signaling pathway, particularly in response to environmental stress (Dittrich et al., 2019; Fidler et al., 2022; Gonzalez-Guzman et al., 2012). Since both Mp*PYL1* and Mp*ANT* are co-expressed in apical cells, it would be interesting to test whether they interact directly and if Mp*ANT* plays a role in ABA signaling in *M. polymorpha*. Additionally, we identified Mp*GRF*, a gene moderately correlated with Mp*ANT* and expressed at higher levels in apical cells. In *A. thaliana*, GRF proteins are linked to proliferative niches in various organs (Liebsch & Palatnik, 2020) and control the expression of *PLT* genes (Rodriguez et al., 2015). Other co-expressed genes were identified in specific regions, such as Mp*ASLBD17* and Mp*RDR5/6-2* in the young sporophyte, and Mp*CHK1*, Mp*LFY*, Mp*BHLH44*, and Mp*DCL3* in the antheridia/archegonia. These findings open new directions to explore the function of Mp*ANT* in other regulatory networks, organs and developmental stages (all gene IDs shown in Figure S5).

### 4.3 Mp*ANT* is required for proper MpPIN1 subcellular localization

In the cells of Mp*ant* gemmae MpPIN1 localization to the plasma membrane is severely affected (Figure 5). This phenotype cannot be explained by the fact that Mp*PIN1* is a transcriptional target of MpANT. Plasma membrane localization of AtPIN1 relies on the phosphorylation status of AtPIN1 in specific residues which are the substrate of the AGC Kinase AtPINOID (Ganguly et al., 2012; Sasayama etal., 2013). MpPIN1 is the sole long PIN protein in *Marchantia* and the residues which are targets of the AGC kinase AtPINOID in AtPIN1 are conserved in MpPIN1 (Fisher et al 2023). Therefore, we speculate that the mis-localization of MpPIN1 in Mp*ant* may be caused by changes in AGC kinase activity, of which there are Marchantia genes that encode for AGC Kinases (Bowman et al 2017; Figure S8). That the promoter regions of each of these putative AGC Kinase encoding genes bears several potential MpANT response elements led us to propose a model where MpANT regulates expression AGC Kinases, and in the absence of MpANT, the down-regulation of AGC Kinase genes would cause MpPIN1 to lack proper phosphorylation and not be localized to the plasma membrane. This is the first report demonstrating Mp*ANT* acting upstream MpPIN1 localization, and further experiments are needed to support this hypothesis.

We found that MpPIN1 is not localized inside the nucleus but instead around it (Figure 6, Figure S7), consistent with the known intracellular trafficking patterns of long AtPINs which also localized around the nucleus, mainly at the endoplasmic reticulum (Barnez et al., 2013). However, this contrasts with a recent study where authors propose that MpPIN1-GFP localizes in the plasma membrane and the nucleus of *Marchantia* epidermal cells (Tang et al., 2024). Mis-localization of MpPIN1 in the M*pant* gemmae (Figure 5 and 6) is correlated with the reduction of rhizoid formation (Figure S2). Therefore, Mp*ANT* is not only important for meristem maintenance, but also for the proper development of rhizoids. Future experiments are needed in order to define the exact mechanism that connects Mp*ANT* with MpAGC-Kinases and MpPIN1 subcellular localization.

## AUTHOR CONTRIBUTIONS

The experimental plan was supervised by Alfredo Cruz-Ramirez. Wet lab and *in silico* experiments were designed by Cruz-Ramírez, Melissa Dipp-Alvarez, and Lorenzo-Manzanarez J. Luis. Wetlab and *in silico* experiments were performed and/or analyzed by Dipp-Alavrez, A, Lorenzo-Manzanarez J. Luis, Espinal-Centeno Annie, Méndez-Alvarez Domingo, Olvera-Martinez Fernando, Flores-Sandoval Eduardo. Leon-Ruiz Jesus, Bowman John L., Arteaga-Vazquez Mario. The manuscript was written by Cruz-Ramirez Alfredo, Dipp-Alavrez, Melissa, Lorenzo-Manzanarez J. Luis and reviewed by all authors.

## ACKNOWLEDGMENTS

LMJL was supported by CONAHCYT-Mexico through a postdoctoral fellowship (488063, 2023-2025). J L-R (CVU 858608) was supported by Consejo Nacional de Humanidades, Ciencia y Tecnología (CONAHCYT) with a PhD Fellowship and CINVESTAV Elisa Acuña grant. MDA was supported by Consejo Nacional de Humanidades, Ciencia y Tecnología (CONAHCYT) with a PhD Fellowship and CINVESTAV Elisa Acuña Grant. EF-S and JLB were funded in part by The Australian Research Council Centre of Excellence for Plant Success in Nature and Agriculture (CE200100015). M.A.A.V was supported by Consejo Nacional de Ciencia y Tecnología (CONACYT) grant A1-S-38383 and UCMEXUS-CONACYT Collaborative Grant CN-20-166.

