## Supplementary material for "The Mp*ANT*-auxin loop modulates *Marchantia polymorpha* development": SupplData

1. Advanced Genomics Unit (LANGEBIO) of CINVESTAV. Km 9.6 Carretera Irapuato-León. CP 36824. Irapuato, Guanajuato, México.

2. School of Biological Sciences, Monash University, Melbourne VIC 3800, Australia.

3. Universidad Veracruzana. Instituto de Biotecnología y Ecología Aplicada (INBIOTECA). Avenida de las Culturas Veracruzanas 101. Colonia Emiliano Zapata. C.P. 91090. Xalapa, Veracruz. Mexico.

+ These authors should both be considered as first authors.

### Supplementary Figures

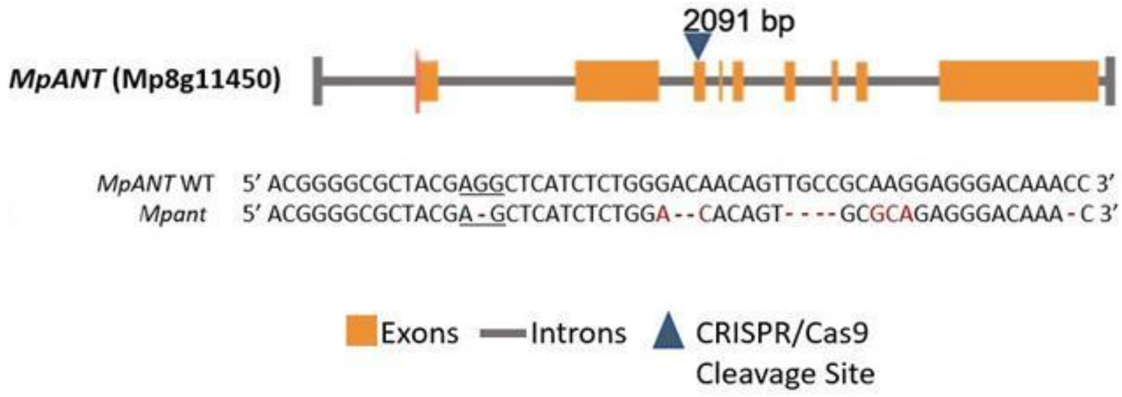


**FIGURE S1** CRISPR-Cas9 edited loss-of-function Mp*ANT* allele. Depiction of the Mp*ANT* (Mp*8g11450*) gene. Cas9 cleavage site targeted with the sgRNA to the third exon (at 2091 bp, blue arrow). Orange boxes represent the exons of Mp*ANT*, while the gray line depicts untranslated regions of the gene. In the nucleotide alignment, the top row shows the Mp*ANT* gene WT sequence. The targeted PAM sequence is underlined. In the bottom row, the sequence of the Mp*ant* mutant line shows deletions and mismatches that causes a premature stop codon which leads to the translation of an incomplete protein.

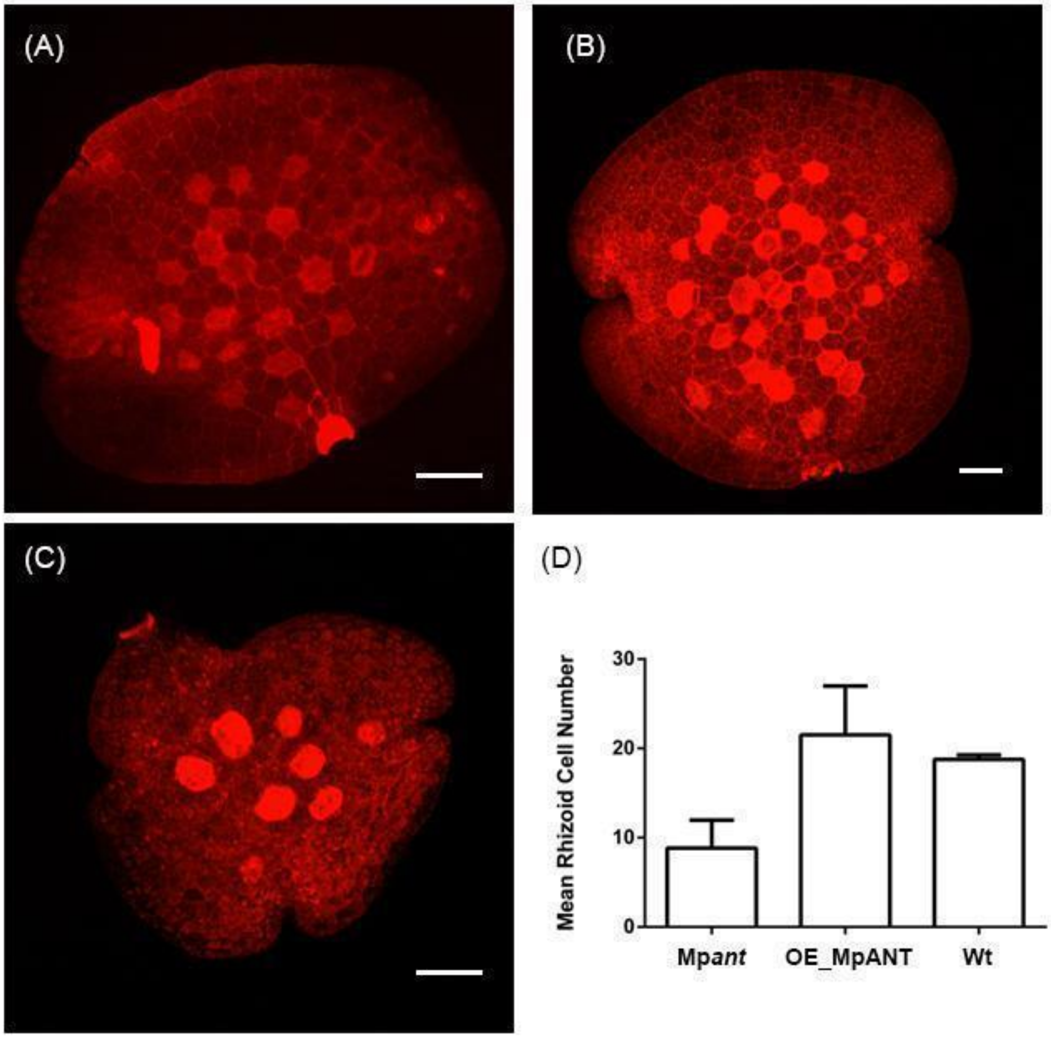


**FIGURE S2. Number of r**hizoid initial cells. Confocal images of 2 day-old gemmae (A) WT, (B) Mp*ANT*$^{OE}$ and (C) Mp*ant*. Data for a minimum of 4 individuals was plotted in (D). Bars in all confocal pictures: 50 uM.

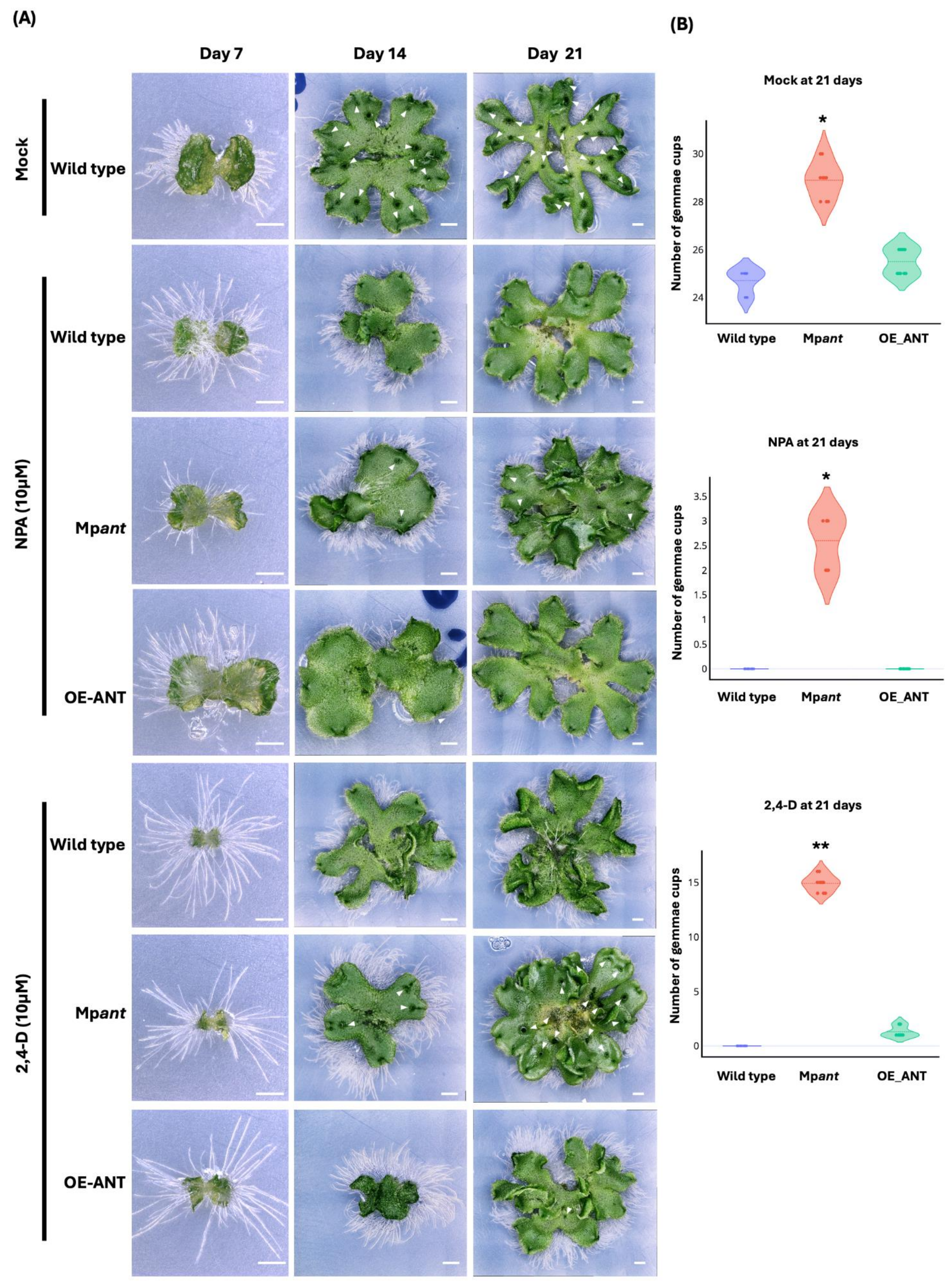


**FIGURE S3.** Phenotypes of Mp*ant*, Mp*ANT*$^{OE}$ and WT plants of 21 days post treatment with NPA (10μM) and 2,4-D (10μM) and controls. (A) NPA and 2,4-D fully inhibits gemmae cup formation in WT and Mp*ANT*$^{OE}$ plants, while in Mp*ant* treated plants the number of gemmae cups drastically decreases when compared to the control. The white arrowhead demarcates gemmae cups. Scale bar=1 cm. (B) Mean of gemmae cups production in plants of the diverse genetic backgrounds grown in Mock, NPA and 2,4-D, respectively. One-way ANOVA followed by Tukey's post hoc test compared to Tak-1(*$P$ <0.05 and **$P$ < 0.01).

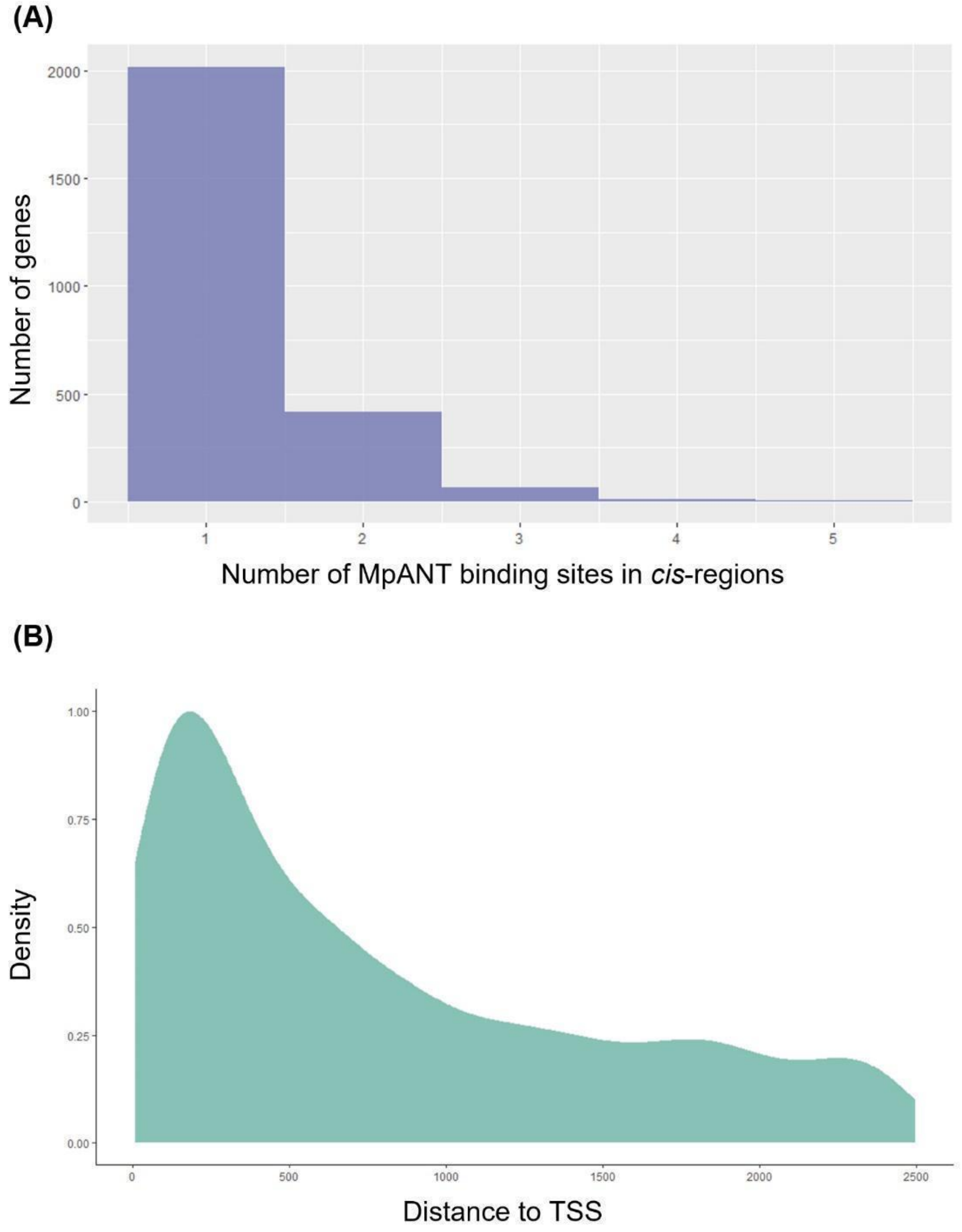


**FIGURE S4**. Number of MpANT binding sites in gene *cis*-regions and their distance to the transcription start site. (A) Histogram showing the number of genes with 1-5 MpANT binding sites in their *cis*-region (2500 bp). (B) Density plot that shows the distribution of MpANT binding sites on the 0-2500 bp region upstream of the transcription start site of putative MpANT target genes.

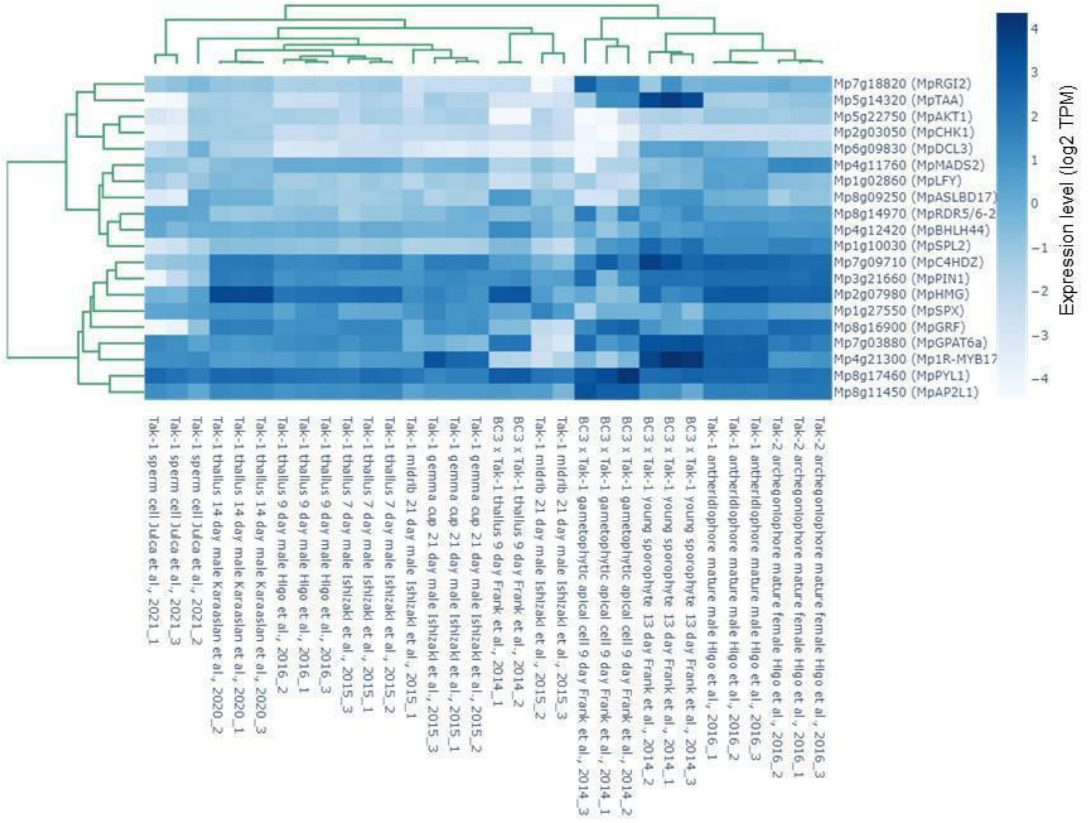


**FIGURE S5.** Hierarchical clustering heatmap of the expression of Mp*ANT* and its putative target genes in different *M. polymorpha* tissues. Rows and columns are re-ordered according to the clustering result. Each line refers to data from a single gene. The gene clustering tree is shown on the left, and the sample or tissue clustering tree appears at the top. The color bar represents the level of gene expression in the sample or tissue (TPM values were log2 transformed), ranging from dark blue (4) to white (-4). Figure and clustering generated using the MarpolBase Expression platform (Kawamura S., et al. 2022).

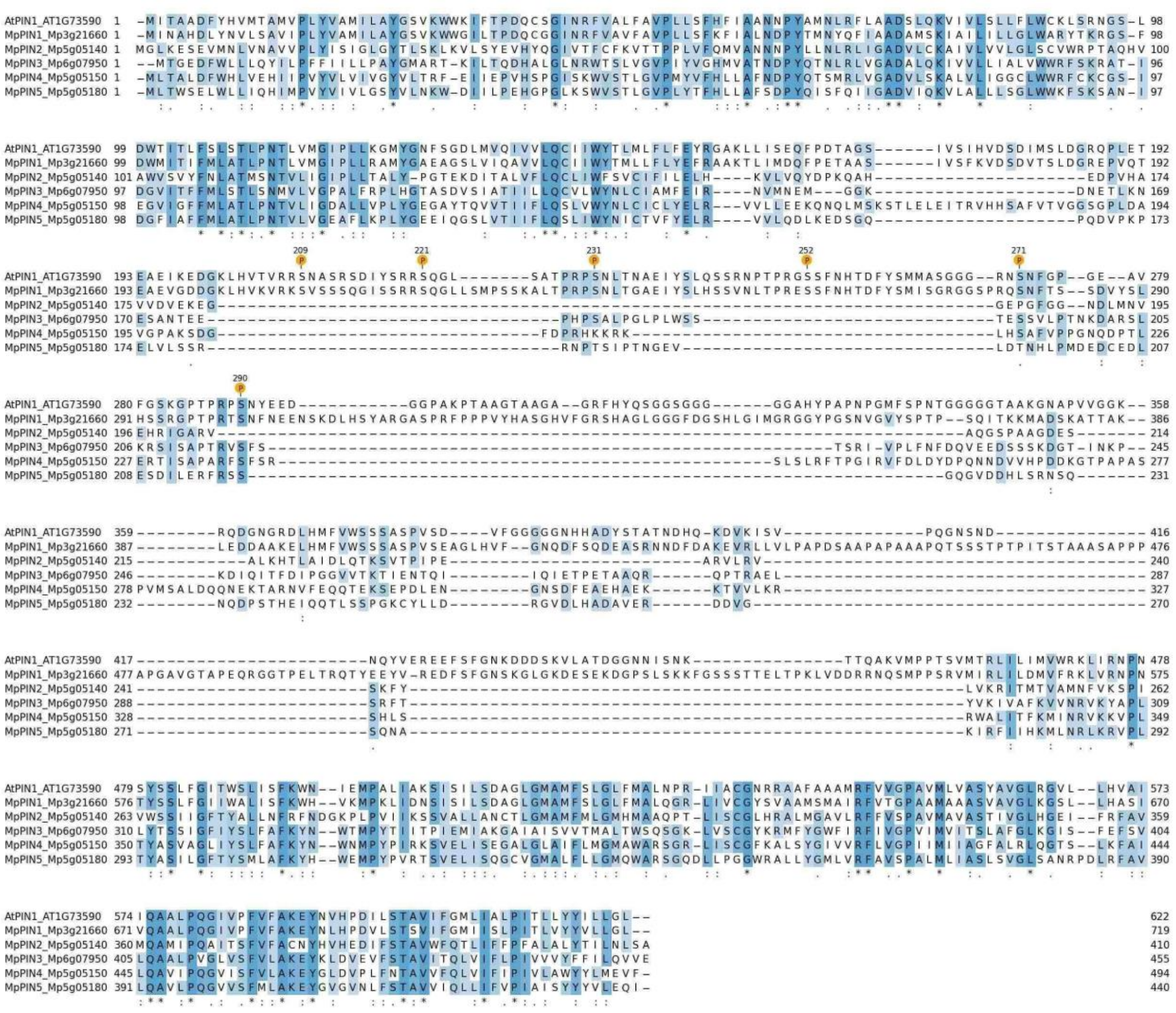

**FIGURE S6.** Alignment of AtPIN1 with the MpPINs from *Marchantia*. The amino acid sequences of the PINs from *Marchantia* were retrieved from https://marchantia.info/. The circles marked with P represent the phosphorylation sites.

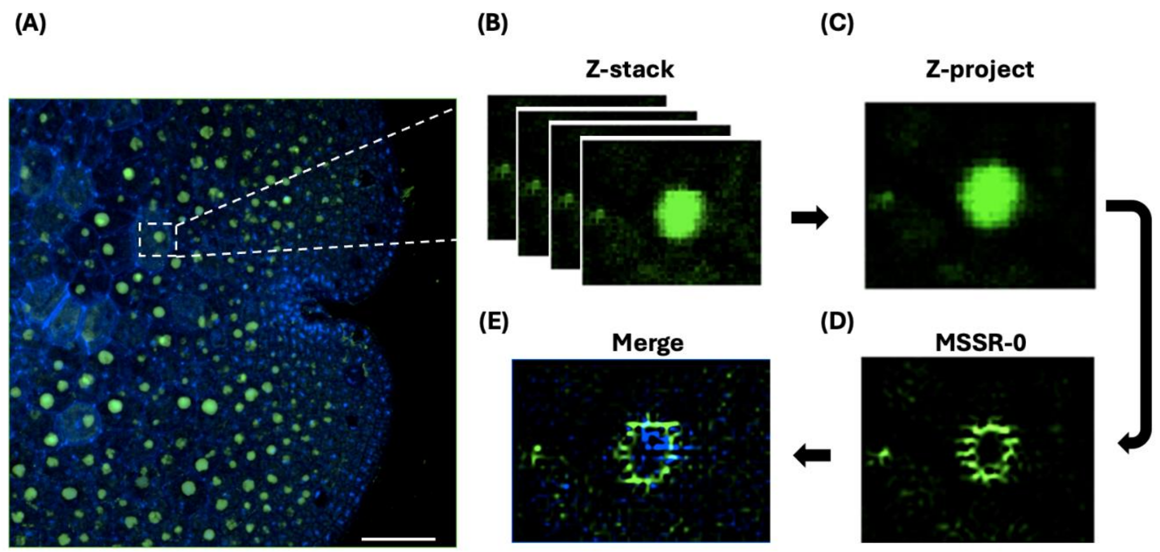


**FIGURE S7**. The localization of MpPIN1 in gemmae cells. A) Apical notch zone and the white dotted lines show nuclei. B) Z-stack of single nuclei. C) Z-project process of nuclei. D) Analysis temporal of MSSR order cero. E) Merge of DAPI signal with new signal of MpPIN1. Scale bars = 50 µm.

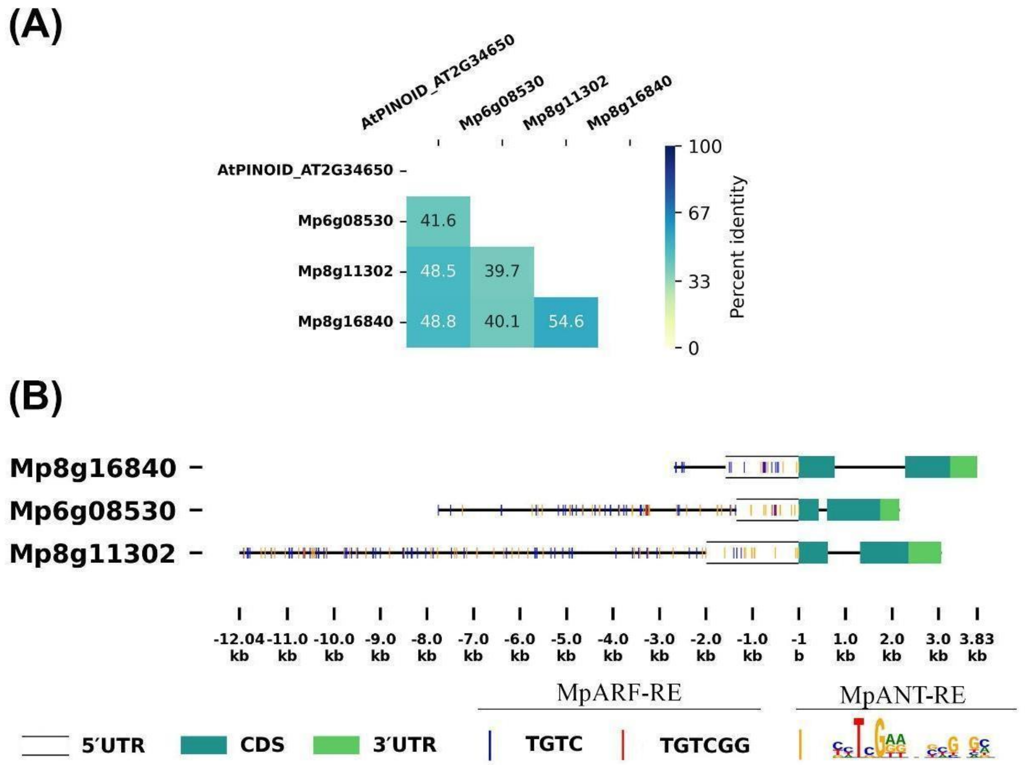

**FIGURE S8**. A) Putative PINOID-like AGC Kinases in *Marchantia*. B) Mp*ARF* and Mp*ANT* cis regulatory elements in the promoter regions of the three AGC Kinases in *Marchantia*.
